# Tau amyloidogenesis begins with a loss of its conformational polymorphism

**DOI:** 10.1101/2020.06.18.158923

**Authors:** María del Carmen Fernández-Ramírez, Rubén Hervás, Margarita Menéndez, Douglas V. Laurents, Mariano Carrión-Vázquez

## Abstract

Knowledge on the molecular bases of early amyloid assembly is fundamental to understand its structure-dysfunction relationship during disease progression. Tauopathies, a well-defined set of neurodegenerative disorders that includes Alzheimer’s disease, are characterized by the pathological amyloid aggregation of tau. However, the underlying molecular mechanisms that trigger tau aggregation and toxicity are poorly understood. Here, using a single-molecule approach, AFM-based single molecule-force spectroscopy (AFM-SMFS), combined with a protein-engineering mechanical protection strategy, we have analyzed the fluctuations of the conformational space of tau during the start of its pathological amyloid assembly. Specifically, we have analyzed the region that includes the four tau microtubule-binding repeats, known to play a key role on tau aggregation. We find that, unlike other amyloid-forming proteins, tau aggregation is accompanied by a decrease of conformational polymorphism, which is driven by amyloid-promoting factors, such as the Δ280K and P301L mutations, linked to Frontotemporal Dementia-17, or by specific chemical conditions. Such perturbations have distinct effects and lead to different tau (aggregate) structures. In addition to providing insight into how tau aggregates in a context dependent manner, these findings may help delve into how protein aggregation-based diseases, like Alzheimer’s, might be treated using monomer fluctuations as a pharmacological target.

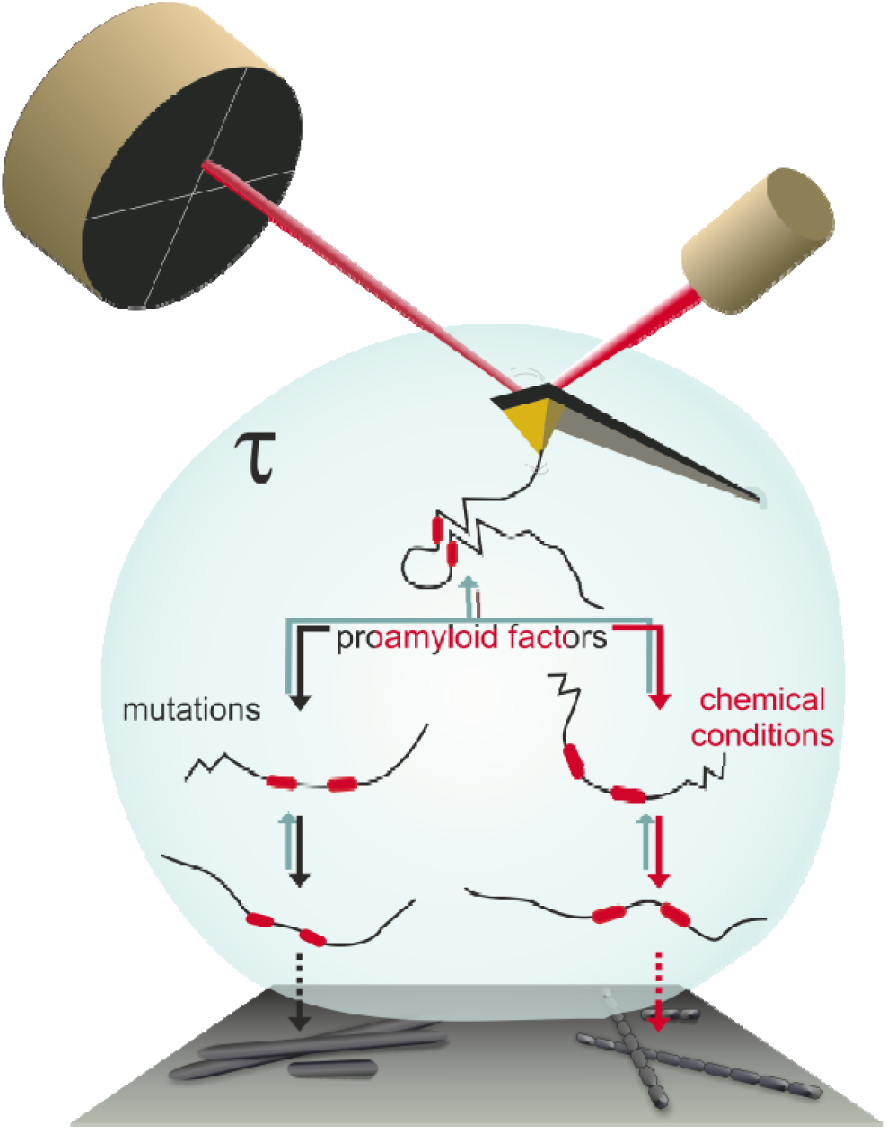

## Introduction

Tau is a human microtubule-associated protein implicated in the regulation of axonal microtubules in the central and peripheral nervous systems (Binder, Frankfurter, and Rebhun 1985; Kadavath et al. 2015). It belongs to the <u>a</u>myloidogenic intrinsically <u>d</u>isordered <u>p</u>rotein (aIDP) class of proteins and in pathological conditions, produces different amyloid aggregates. These aggregates are associated with neurodegenerative disorders called tauopathies, including Alzheimer’s disease, Frontotemporal Dementia and Parkinsonism Linked to Chromosome-17 (FTDP-17) (Avila et al. 2004). The full-length tau isoform contains 441 residues, which can be divided in two domains: the N-terminal region, which is the projection domain comprising about two thirds of the molecule; and the C-terminal region, which contains the <u>M</u>icrotubule <u>B</u>inding <u>D</u>omain (MBD). The high solubility of MBD is due by its positively charged residues in physiological conditions, which also promote the binding to the negatively charged microtubules. Net charge of constructs including only the MBD with either three (K19 fragment) or four (K18 fragment) pseudo repeats is +7 and +10, respectively, and their isoelectric point is 10.5 (Jeganathan et al. 2008). Both fragments are commonly used *in vitro* to study the aggregation properties of tau (Jeganathan et al. 2008; Shammas et al. 2015). Furthermore, they undergo liquid/liquid <u>p</u>hase <u>s</u>eparation (LLPS) (Ambadipudi et al. 2017), which promotes its aggregation (Wegmann et al. 2018) forming the typical <u>p</u>aired <u>h</u>elical filaments (PHF) of tau while excluding the flanking regions, known to slow down tau amyloid aggregation (Wille et al. 1992). The K18 fragment, which spans residues 244-368, shows higher ability for demixing (Ambadipudi et al. 2017) and aggregation (Wille et al. 1992; Daebel et al. 2012) than K19, and includes the two six-residue segments ^275^VQIINK^280^ and ^306^VQIVYK^311^, which are drivers for amyloid formation (von Bergen et al. 2000; 2001) **(Figure 1A)**.

**Figure 1.**
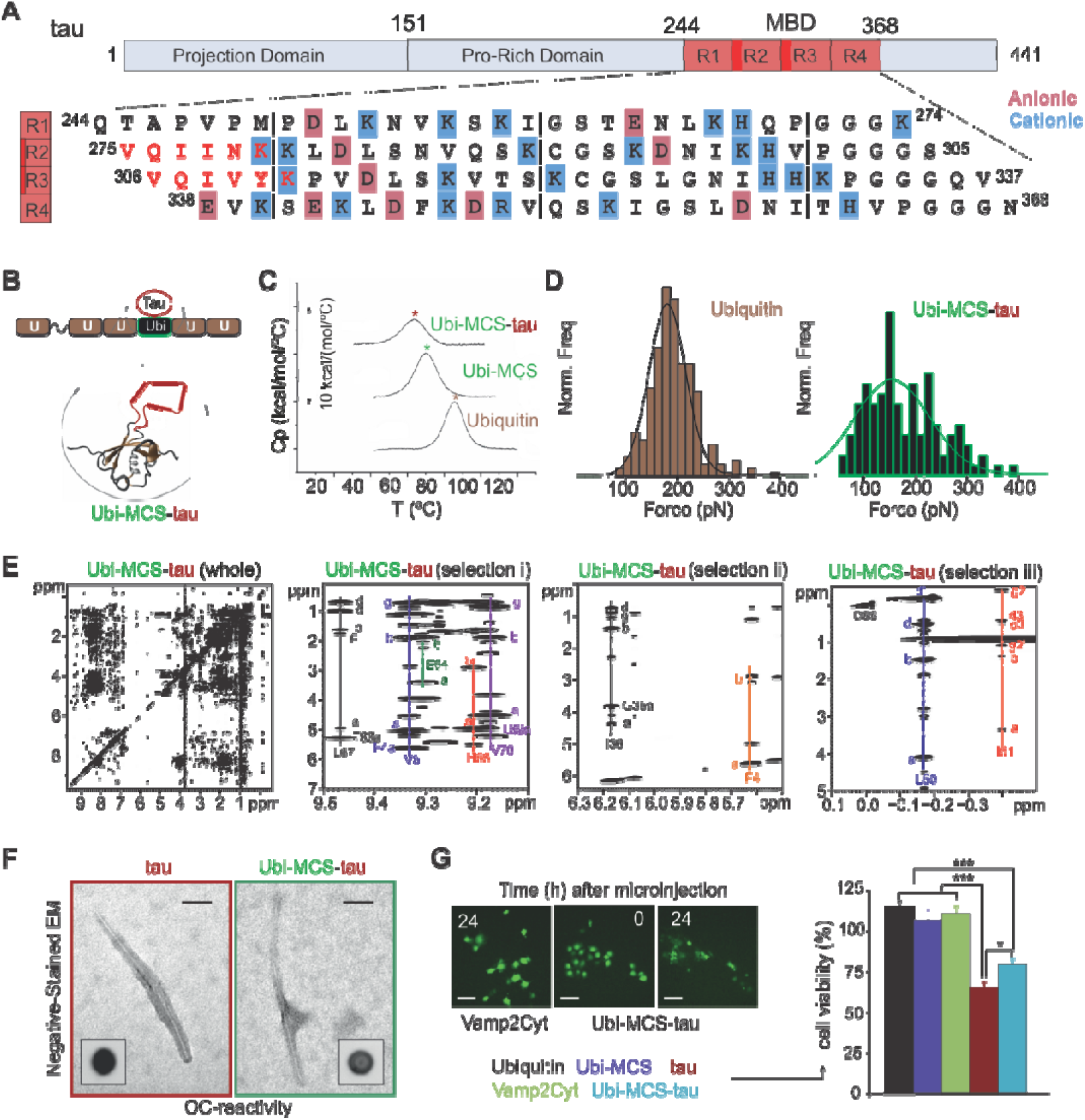
Tau protein and validation of the carrier-guest strategy. **A)** Scheme of the longest tau isoform and the sequence of the K18 fragment used in this study. Charged residues and pro-aggregant six-residue segments are highlighted. **B)** Schematic cartoon of the pFS-2 vector using a Ubi as a carrier and hosting the tau fragment K18. The fusion protein Ubi-tau is highlighted. **C)** Thermal and **D)** mechanical unfolding analyses to test the integrity of Ubiquitin in the Ubi-tau construction. Upon tau insertion, Ubi is slightly destabilized, as it has been described for other carrier-guest constructions in previous studies (Hervas et al. 2012), which still allows AFM-SMFS analysis. **E)** Complete (i) and selected regions (ii, iii and iv) of Ubi-tau NOESY spectrum ^1^HN region. The crosspeaks from certain deshielded ^1^HN of natively folded ubiquitin to alpha (a), beta (b), gamma (g) and delta (d) protons are labeled. Crosspeaks from ^1^HN to the ^1^H alpha of the preceding residue are labeled in three cases. iii) The crosspeaks between the 1HN of I36 (6.17 ppm) and its ^1^H alpha, the ^1^H alpha of G35 and the side chain protons of I36 are labeled. The ^1^H alpha of F4 (on the diagonal) and the crosspeaks to its beta protons are also labeled. iv) Crosspeaks arising from the shielded delta ^1^H of L50 and the shielded gamma methylene ^1^H of I61 are shown. For I61, g2’ refers to the other gamma methylene proton, d3 is the delta methyl protons and g3 is the gamma methyl protons (Di Stefano and Wand 1987). DSS (3-(Trimethylsilyl)-1-propanesulfonic acid) is used as chemical shift reference. **F)** After 7 days of incubation at 37°C, both tau, isolated and inside the Ubi carrier, form amyloid-like fibers, which are recognized by the anti-amyloid OC antibody. Scale bar: 100 nm. **G)** Tau microinjection in the COS-7 cell line. Left panel: fluorescence micrographs of microinjected COS-7 cells. Scale bars: 100 *μ*m. Right panel: cell viability, after 24h post-microinjection. Only tau and Ubi-tau proteins cause toxicity. Data are represented as the mean ± SEM: \**p*<0.05; \*\*\**p*<0.005 (red asterisks) tau and Ubi-tau *versus* remaining samples. Ubi, Ubi-MCS, and the cytoplasmic region of VAMP2 are used as controls of folded and disordered proteins, respectively.

Tau has been extensively studied by structural (Jeganathan et al. 2008; von Bergen et al. 2000; Mukrasch et al. 2009; Fitzpatrick et al. 2017; Falcon et al. 2018; 2019; W. Zhang et al. 2020), single-molecule (SM) (Wegmann et al. 2011; Manger et al. 2017) and computational approaches (Larini et al. 2013). In summary, these reports show that monomeric tau adopts a random coil conformation that evolves, through a progressive *β*-structure acquisition, to form the amyloid state. The amyloid fibrils assembled *in vitro* are polymorphic (Wille et al. 1992; Xu et al. 2010; W. Zhang et al. 2019), while disease-specific structures for tau extracted from patient brains of AD (Fitzpatrick et al. 2017), Pick’s disease (Falcon et al. 2018), chronic traumatic encephalopathy (Falcon et al. 2019) and corticobasal degeneration (W. Zhang et al. 2020) have been also reported. An additional layer of complexity is added with the LLPS formation (Ambadipudi et al. 2017). Although the amyloid state of tau is being widely studied, the conformational landscape of monomeric tau and how it is perturbed during amyloid assembly is still under debate. As a result, no therapeutic targets at the start of the amyloid assembly pathway of tau have been identified.

In order to study the conformational landscape of monomeric tau at single-molecule resolution, we have used <u>A</u>tomic <u>F</u>orce <u>M</u>icroscopy (AFM) in the <u>S</u>ingle-<u>M</u>olecule <u>F</u>orce <u>S</u>pectroscopy (SMFS) mode, together with a mechanical protection strategy for unequivocal single-molecule identification (Oroz, Hervas, and Carrion-Vazquez 2012). This approach has been previously used to analyze the start of the amyloid assembly pathway of pathological (Hervas et al. 2012) and functional (Hervas et al. 2012; 2016) amyloid-forming proteins. Using this technique, the intramolecular interactions stabilizing a protein conformation can be mapped and measured (Carrion-Vazquez et al. 2000). Unlike other amyloid-forming proteins (Hervas et al. 2012), we found a decrease of tau conformational polymorphism associated with its amyloid aggregation. Furthermore, the analysis of the mechanical properties indicates that distinct pro-aggregative factors lead to different monomer conformations as well as final amyloid structures. These results point to nascent conformations in the monomer as key therapeutic targets given their appearance at the beginning of the amyloidogenic cascade.

## Results and discussion

### Validation of the chosen strategy to study tau nanomechanics

To analyze the nanomechanics of monomeric tau, we selected the K18 tau fragment (called “tau” for short through the text), cloned in the pFS-2 vector (Oroz, Hervas, and Carrion-Vazquez 2012). This vector, based on the carrier-guest strategy, was developed for the AFM-SMFS analysis of disordered proteins, particularly aIDPs. In this approach, tau is mechanically protected by a carrier protein, consisting of a modified ubiquitin that bears a <u>m</u>ulti<u>c</u>loning <u>s</u>ite (Ubi-MCS) **(Figure 1B)**. By cloning tau in the artificial MCS, located in an Ubi loop after the mechanical clamp, we were able to insert tau into the ubiquitin fold. Hence, in a typical AFM-SMFS experiment, tau will only be unraveled after the mechanical unfolding of the carrier.

Before carrying out the AFM-SMFS measurements, we tested on the isolated Ubi-tau protein whether the properties of the engineered tau and the Ubi carrier are preserved in Ubi-tau fusion protein by performing standard structural, calorimetric, and fibrillogenic assays **(Figure 1C-G)**. The structural integrity of the carrier Ubi was measured first by thermal and mechanical denaturation, since thermal and mechanical stabilities are not necessarily correlated (Carrion-Vazquez et al. 2000). <u>D</u>ifferential <u>s</u>canning <u>c</u>alorimetry (DSC) showed that upon MCS insertion, a drop in the Ubi denaturation temperature (Tm), from 85.5 °C to 70.5 °C, was observed. Upon tau insertion the Tm decreased further to 63.2 °C **(Figure 1C)**. AFM-SMFS experiments on pFS-Ubi-tau allowed us to determine the mechanostability of Ubi and Ubi-MCS carrying tau, by measuring the unfolding force *F*_u_ of Ubi single-molecule markers (n=282) and Ubi-MCS carrier modules (n=112), respectively. As expected from previous results, the average *F*_u_ for Ubi-MCS-tau was lower, and with more scattered values, than that of native ubiquitin (182 ± 50 pN *versus* 157 ± 104 pN) **(Figure 1D)**. This thermal and mechanical destabilization is comparable to that reported for other amyloid insertions (Hervas et al. 2012), which indicates that the Ubi carrier is suitable for the nanomechanical analysis of tau.

In addition to this, we performed an analysis of the structure of Ubi-tau by NOESY NMR. The spectrum of Ubi-tau showed chemical shifts that are in close agreement with those obtained for the isolated WT Ubi under similar conditions (Di Stefano and Wand 1987). Moreover, no indication of measurable contacts between the carrier and tau were detected **(Figure 1E)**. Fibrillogenic tests were also performed to determine whether the amyloidogenic behavior of tau remained unaltered in the fusion protein. After seven days of incubation at 37°C, Ubi-tau, like the isolated tau, formed unbranched fibers visualized by EM, which were reactive against the conformational anti-amyloid OC antibody (Kayed et al. 2007) **(Figure 1F)**. Finally, since the goal of this study is to identify early structural features that may lead to cytotoxicity, the toxic effects of both tau and Ubi-tau were measured by microinjecting them in single COS-7 cells after 7 days of incubation at 37°C **(Figure 1G)**. Together, these results show that the Ubi carrier retains its native 3D structure unaltered upon tau insertion, while tau conserves their aggregative and toxic capabilities in the fusion protein, which shows that the mechanical protection strategy of tau is suitable for its AFM-SMFS nanomechanical analysis.

### Tau conformational polymorphism monitored by SMFS

We used SMFS to analyze the nanomechanics of tau, expressed as a fusion polyprotein in pFS-2 (Oroz, Hervas, and Carrion-Vazquez 2012) **(Figure 2A)**, and followed a strict set of criteria for selecting valid recordings (see Materials and Methods section). Using the length-clamp mode, two basic parameters are directly obtained: *i)* the mechanical stability of the resistance barriers, *F*_u_, which corresponds to the height of the force peaks in a force-extension trace; and *ii)* the length of the force-hidden region, increase in contour length Δ*L*_*c*_, as measured by fitting the Worm-like chain model of polymer elasticity to the force-extension trace **(Figure 2B)**. Tau SMFS analysis showed that the WT form exhibited different populations of conformers. Thus, we found that 49% of molecules generate Non-Mechanostable (NM) events, where there are no SMFS-detectable conformations (*e.i*., no detectable force peaks, *F*_u_≤20 pN, the force detection limit of SMFS at a pulling speed of 0.4 nm/ms). In the NM recordings the Δ*L*_*c*_ signal from the entire tau fragment appears immediately after the unfolding of the carrier **(Figure 2B)**. These events are typically generated by conformers with random coil or isolated *α*-helix structures (Oberhauser and Carrion-Vazquez 2008). We also found that 51% of the molecules belongs to a mechanostable (M) population (*i.e*., with recordings carrying force peaks, *F*_u_>20 pN), which are likely originated from the unravelling of *β*-structured conformers (Oberhauser and Carrion-Vazquez 2008) or coil-coiled structures (Brown et al. 2007; Goktas et al. 2018; Bornschlogl and Rief 2008; Schwaiger et al. 2002). The M population showed one or several detectable force peaks, originated from the rupture of their intramolecular interactions, with a variety of Δ*L*_c_ values **(Figure 2B**), which indicates a rich conformational polymorphism of the WT tau. Computational studies have previously predicted this conformational diversity, in particular, for fragments containing the hexapeptides ^275^VQIINK^280^ or ^306^VQIVYK^311^ contained in the K18 fragment that we have analyzed here (Larini et al. 2013; D. Chen et al. 2019). Inside the M class we found that 2.6% of the molecules form rare hyper-Mechanostable (hM) conformers (operationally defined as *F*_u_≥400 pN) **(Figure 2B)**, a common feature shared by all the aIDPs, both pathological and functional, analyzed so far (Hervas et al. 2012; 2016). The hM conformers were initially proposed to be involved in cell toxicity by blocking the proteasome and/or other protein degradation systems (Hervas et al. 2012; 2016). However, this hypothesis should be reformulated based on both the fact that functional amyloids also show these highly-stable conformers (Hervas et al. 2012; 2016) and on the consideration that the AFM pulling geometry *in vitro* is different to that of the proteasome or the chaperons *in vivo* (Wojciechowski et al. 2014).

**Figure 2.**
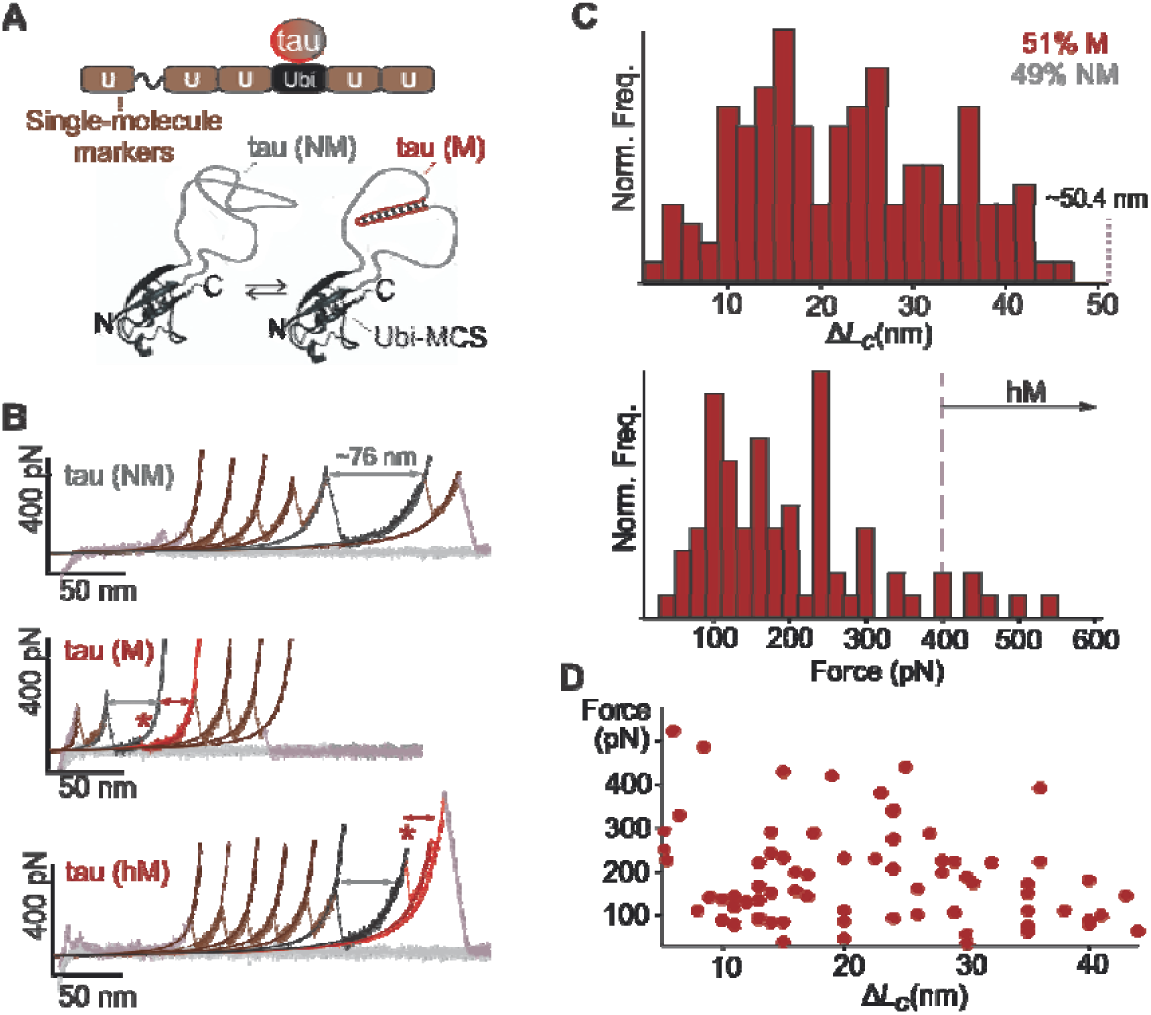
Tau nanomechanics. **A)** Schematic representation of the pFS-tau polyprotein which illustrates the color code used for the representation of the SMFS data. Inside the ubiquitin carrier, tau can freely fluctuate between non-mechanostable, NM (gray), and mechanostable, M (red) conformations. **B)** From top to bottom, examples of NM, M and hM recordings are shown. Brown traces correspond to ubiquitins, the primary controls for selecting single molecules. The NM conformer (top trace) shows non-detectable stability mechanical barriers and is only detected once Ubi carrier is mechanically unfolded. The middle and bottom recordings show M conformations with different degree of mechanical stability (red asterisks indicates the first mechanical barrier), including hM events (bottom trace). Dark traces and gray arrows indicate NM signal coming from Ubi-MCS and tau stretching while red arrows indicate acquired tau structures that form M events. **C)** Structural polymorphism of tau. Δ*L*_*c*_ (top panel) and *F*_u_ (bottom panel) histograms the pFS-tau polyprotein (n=114) indicate that tau forms intramolecular interactions all along its sequence, which are heterogeneous in mechanostability, including hM conformers (with *F*_u_ over 400 pN). **D)** Scatter plots show that there is no correlation between *F* and Δ*L*_c_ values, which indicates that WT tau does not form preferential conformers.

### Modulators of tau nanomechanics: pathological mutations and buffer composition

Several reports found that tau amyloid assembly can be promoted or prevented *in vitro*. In order to understand whether this regulation is exerted at the monomer level and to further explore the molecular details of the early tau amyloid pathway, we have examined a number of factors influencing the process. First, the effect of the anti-amyloidogenic polyQ-binding peptide QBP1 (Nagai et al. 2007; Ramos-Martin et al. 2014) on tau nanomechanics was tested. Similar to the results obtained with A*β*, and contrary to other aIDPs such as polyQs, *α*-synuclein, Sup35NM (Hervas et al. 2012), TDP-43 (Mompean et al. 2019), Orb2 (Hervas et al. 2016), *Ap*CPEB (Hervás et al. 2020) and hCPEB3 (Ramírez de Mingo et al. 2020), QBP1 did not produce a significant effect on the conformational polymorphism of monomeric tau or in its aggregation capabilities **(Figure S1)**. QBP1 was initially developed to inhibit the aggregation the expanded polyQ tracts and was found to prevent the toxic *β*-sheet conformational transition in the monomer (Nagai et al. 2007), a plausible species that would act as a nucleus for aggregation (Hervas et al. 2020) in a “nucleation growth” model (S. Chen, Ferrone, and Wetzel 2002; Hervas et al. 2020). The lack of inhibitor action on tau could be due to incompatibilities with either the physico-chemical properties of the protein, such as not having polyQ segments, or with the molecular mechanism underlying its amyloidogenic process, or to both.

Second, two pro-amyloidogenic mutations, and different buffer conditions, were analyzed as additional possible monomer modulators. The selected pathological mutations, Δ280K and P301L have a causal role in hereditary <u>F</u>rontotemporal <u>D</u>ementia with <u>P</u>arkinsonism-<u>17</u> (FTDP-17) and they have been shown to increase tau amyloid formation, *in vitro* (S Barghorn et al. 2000; von Bergen et al. 2001). As for the buffers, according to previous findings (Jeganathan et al. 2008; Jebarupa et al. 2018), we selected three different buffer conditions, namely, *i)* antiamyloidogenic (PBS, pH 7.4), *ii)* neutral (10 mM TrisHCl, pH 7.4) and *iii)* proamyloidogenic (10 mM TrisHCl, pH 10.0, 3:1 heparin). As we did for WT tau, the Ubi carrier fused with tau harboring Δ280K and P301L mutations were validated by far-UV and ^1^H NMR prior the SMFS analysis **(Figure S2)**.

Then we quantified by SMFS the M and NM conformers in the same conditions used for the WT tau (the neutral condition was used for the initial characterization of tau**; Figure 2)**. Using this buffer, 117 valid recordings for Δ280K_tau and 151 for P301L_tau were selected and analyzed, revealing 63% and 78% NM events, respectively. For the WT tau, 112 curves were analyzed in the anti-amyloidogenic buffer condition and 102 in the proamyloidogenic one, finding 44% and 63% NM events, respectively. The application of χ^2^ tests of independence allowed us to conclude that both variables, “mutations” (*p*<0.001) and “buffer” (*p*<0.005), significantly modulate the space of possible acquired tau conformations at the monomer level **(Figure 3)**. Thus, both variables were used in the subsequent analysis. In addition to this, the abundance of M events and the number of force peaks/M event were quantified. The M events are assumed to arise from conformers that have acquired secondary or tertiary structures that lead to the formation of a more compact conformation (Lambrughi et al. 2012). Contrary to the findings in other neurotoxic proteins (Hervas et al. 2012), this analysis revealed that the conditions that discourage tau aggregation are associated with a higher number of both M events and force peaks/M event (Hervas et al. 2012). By contrast, when tau is perturbed by proaggregation mutations or is kept in the proamyloidogenic buffer, preferentially adopted NM conformations. These data indicate that amyloid aggregation of tau is associated to unstructured and consequently expanded monomers, suggesting an alternative assembly pathway **(Figure 3)**. Our results are supported by computational studies describing a shift in tau peptides carrying Δ280K or P301L towards more expanded conformations (Larini et al. 2013; D. Chen et al. 2019).

**Figure 3.**
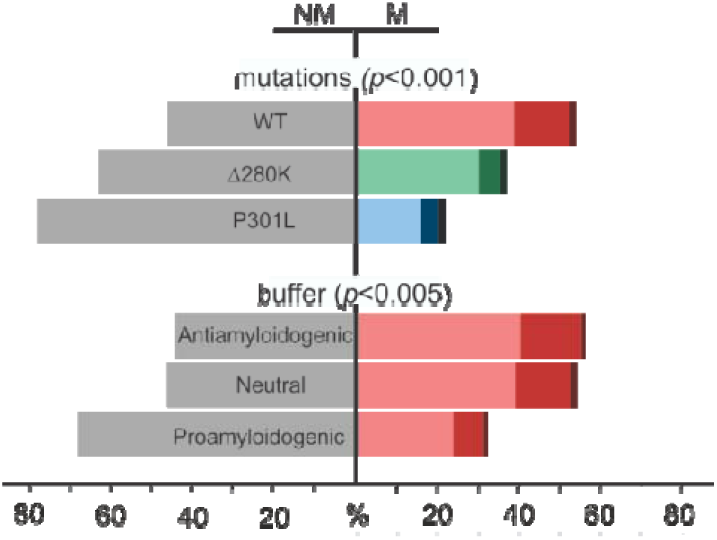
Quantification of tau mechanical conformers under different conditions. The effect of “mutations” and “buffer” on the NM/M ratios is statistically significant. The frequency of the unstructured population (NM) appears in gray, while the structured group (M) appears in reddish color, which is divided in subpopulations that show 1, 2 or 3 force peaks (from low to high color intensity, respectively). **Top panel:** Comparison among WT, Δ280K and P301L tau sequences analyzed in a neutral buffer. The number of NM events increases in the mutated tau, while the number of structures *per* molecule decreases. **Bottom panel:** behavior of tau WT in conditions with different propensity to form amyloid. A similar trend to reduce intramolecular interactions is shown in proaggregation environments.

The proaggregation effect of heparin in tau has been widely studied (Ramachandran and Udgaonkar 2011; Zhu et al. 2010; Eschmann et al. 2017). Heparin, along with a specific buffer composition, drives the expansion of ^275^VQIINK^280^ and ^306^VQIVYK^311^ segments as we have found in our SMFS analysis from heparin-containing proamyloidogenic buffer **(Figure 3)**. Although those studies do not distinguish whether this phenomena occurs before or after the formation of soluble aggregates (Eschmann et al. 2017), our data suggest that it happens at the monomer level.

It has been previously found that the reduction of cysteines in tau assists tau aggregation *in vitro* (Stefan Barghorn, Biernat, and Mandelkow 2005), while cysteines oxidation is the mechanism used by the compound TRx0237, which is being clinically tested as a drug acting on tau amyloidogenesis (Bakota and Brandt 2016). The reduction of the intramolecular interactions might cause the exposure of the two amyloid motifs ^275^VQIINK^280^ and ^306^VQIVYK^311^, which are drivers in tau aggregation (von Bergen et al. 2000; 2001). These hexapeptides were shown to form steric zippers (Sawaya et al. 2007; Seidler et al. 2018). Furthermore, the dimeric tau has been proposed to form the minimal aggregated species (Congdon et al. 2008; Patterson et al. 2011; Kim et al. 2015). Based on all this, we propose a model were two expanded tau chains with their amyloid-forming motifs exposed could first establish intermolecular interactions in an early dimeric state (see **Figure S5**). These species would undergo the needed structural reorganization that would generate the *β*-structured aggregation nuclei, compatible with the conformational conversion model previously proposed for tau amyloid aggregation (Eschmann et al. 2017).

### Analysis of the acquired tau conformations

So far our results point out to the unfolded monomer as a key species in tau amyloidogenesis, however it remains to know how different conditions perturb the formation of the amyloid-competent monomer that triggers the assembly. To gain insight into this question, the tau structures that remained after each perturbation were further analyzed. To this end, we have focused on the first mechanical barrier observed in an M event upon tau unraveling **(Figure 2B)**, examining mechanostability (*F*_u_) and length (Δ*L*_*c*_) of the structures associated with the barrier, in every experimental condition tested, separately and in relation with the other populations generated. No preferential conformations were formed in any of the conditions assayed, as suggested by the Δ*L*_*c*_ *vs. F*_u_ scatter plots **(Figure S3)**. Isolated *F*_u_, fitting to a log-normal distribution **(Figure 4A-C)**, were transformed into a Gaussian distribution by calculating the logarithm of each value for later comparison of the populations by one-way ANOVA and *post-hoc* Tukey test **(Figure S4)**. In this analysis, no *F*_u_ differences were found among the structures acquired by the different tau variants (WT, Δ280K and P301L). However, when WT tau in the neutral buffer is compared with the protein in the proamyloidogenic buffer, we find that the *F*_u_ of its first mechanical barrier is considerably lower, indicating an overall mechanical destabilization of the remaining structures **(Figure 4D)**. In principle, we could expect that less mechanostable conformers would be more readily unraveled, being on-pathway to the RC species, *i.e*. the amyloid-competed monomer postulated here.

**Figure 4.**
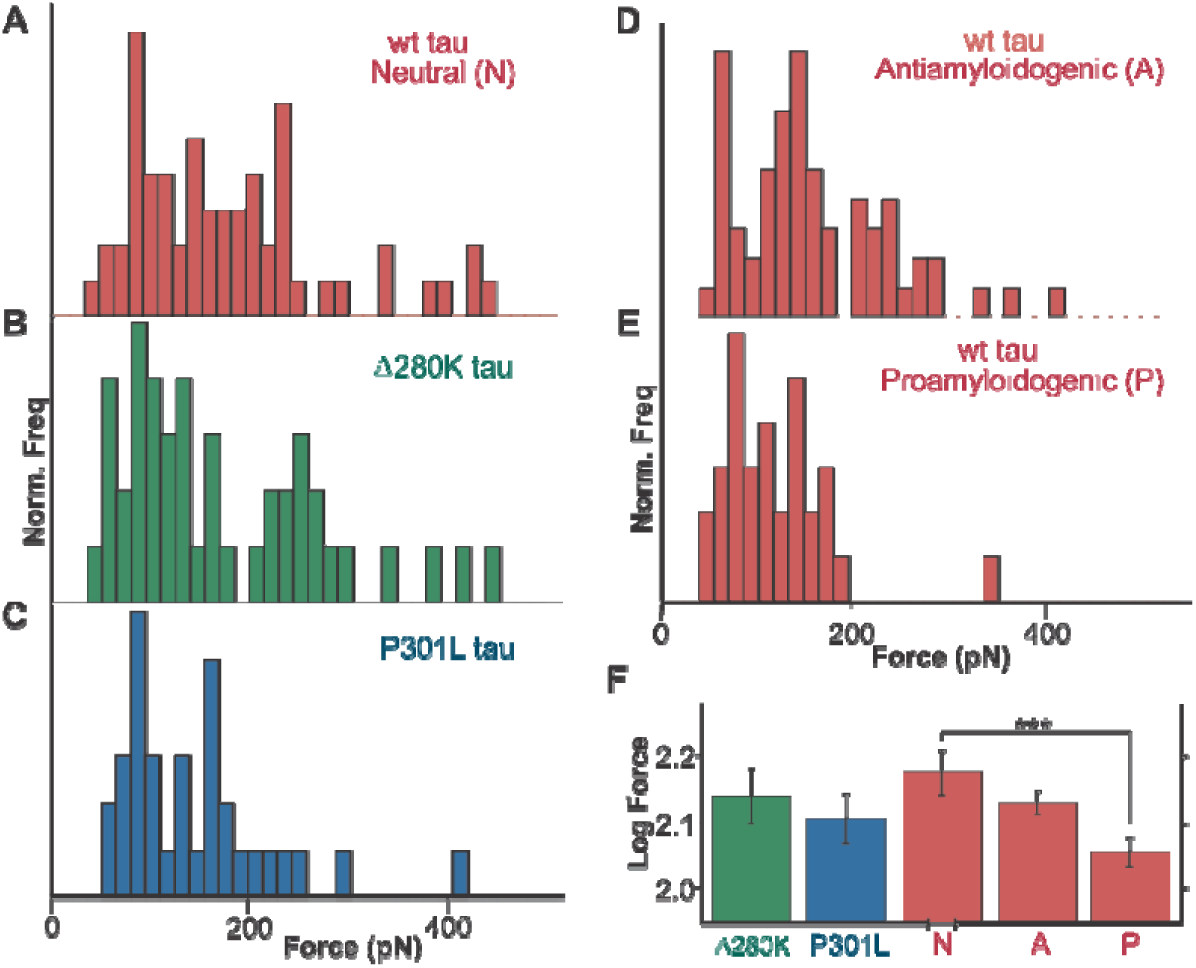
Mutations and buffer effects in the *F*_u_ of the first mechanical barrier of tau conformers. **A)** Tau WT, **B)** Δ280K and **C**) P301L studied in the neutral buffer are represented in different red, green and blue colored panels, respectively. **A, D** and **E)** tau WT studied in different aggregation conditions, represented in red. **F)** Mean, SEM and comparative analysis of the normalized populations obtained under the effect of each variable. A drop in mechanostability is observed in the structures acquired by tau WT in the proamyloidogenic condition. No effect is observed in the *F*_u_ of the Δ280K and P301L mutations.

Based on the theory of the phase diagram of a polymer, subtle changes in the charge of a polypeptide could switch the state of an aIDP (Bergeron-Sandoval, Safaee, and Michnick 2016). In the proamyloidogenic buffer, the pH coincides with K18 isoelectric point and the negatively-charged heparin is present. Thus, the dramatic decrease of the positive net charge causes a reduction **(Figure 3)** and destabilization **(Figure 4D)** of the intramolecular structures. A reduction of the net charge might also be achieved by negatively charged elements also found in the cellular context, such as ARNs (X. Zhang et al. 2017), phosphor-groups (Ambadipudi et al. 2017) or tubulin (Hernández-Vega et al. 2017), promoting a condensed state that may preceded tau amyloid aggregation (Vanderweyde et al. 2016; Wegmann et al. 2018). Our data suggest that the rich conformational space explored by tau should be tightly regulated in neurons in order to fulfill its physiological functions. Misregulation caused by pathological mutations or hyper-phosphorylation would generate an unbalanced increase in the unfolded population of conformers. Based on the blob model for polymer solutions (Uematsu, Svanberg, and Jacobsson 2005), tau might (evolve?) from a collapsed globule, to an expanded globule, and to rigid-rod-like structures (Bergeron-Sandoval, Safaee, and Michnick 2016), *i.e.*, the formation of insoluble fibrils **(Figure S5)**. Supporting this idea, Δ280K and P301L mutants, whose unstructured NM populations are higher than in the WT variant **(Figure 3)**, do not require being phosphorylated to drive LLPS formation, while the WT form does (Wegmann et al. 2018).

Finally, we analyzed the Δ*L*_*c*_, *i.e.* length of the disordered region prior to the first mechanical barrier, considering the 50.4 nm length of the 126 amino acid residues of K18, with an estimate of 0.4 nm/aa (Ainavarapu et al. 2007). Δ*L*_*c*_ values were found to be distributed from 0 to 50.4 nm **(Figure 5A-E)**, which indicates that the first structure can be formed all along the molecule. Except for the anti-amyloidogenic condition, the median of the scattered populations corresponds approximately to the middle point of the tau sequence **(Figure 5F)**, which indicates no preferential structure formation at any specific region. A sub-population of these conformers might be the origin of different aggregation nuclei, accounting for the heterogeneous structures and/or fiber morphologies described for tau aggregates both *in vitro* (Xu et al. 2010; Sanders et al. 2014) and in tau-associated diseases, like Alzheimer’s (Fitzpatrick et al. 2017), Pick’s (Falcon et al. 2018), chronic traumatic encephalopathy (Falcon et al. 2019) and corticobasal degeneration (W. Zhang et al. 2020; Arakhamia et al. 2020). The analysis of the differences between populations revealed that the mutations did not produce any effect in the length of the acquired structures, while a slight perturbation was found between those structures generated in non-promoting amyloid buffers **(Figure 5D)**. The antiamyloidogenic buffer has a high salt concentration, which is known to discourage LLPS formation (X. Zhang et al. 2017). The analysis of the differences between populations **(Figure 5C, D)** revealed an increase in conformers whose first mechanical barrier is located after the unraveling of around 75 aa (≈30 nm /0.4 nm per aa), suggesting that ≈51 aa (126-75) remain hidden to the solvent. Hydrophobic interactions have been described as the likely responsible for an initial collapse that brings the monomers together during LLPS (Schmidt and Gorlich 2016). The hydrophobic hexapeptides ^275^VQIINK^280^ and ^306^VQIVYK^311^, present in the K18 fragment here employed, are separated by 36 residues, enabling a possible interaction from the flanking regions to hide the 51 residue-long region. Thus, those segments would first drive for LLPS and then latter promote amyloid aggregation **(Figure S5).**

**Figure 5.**
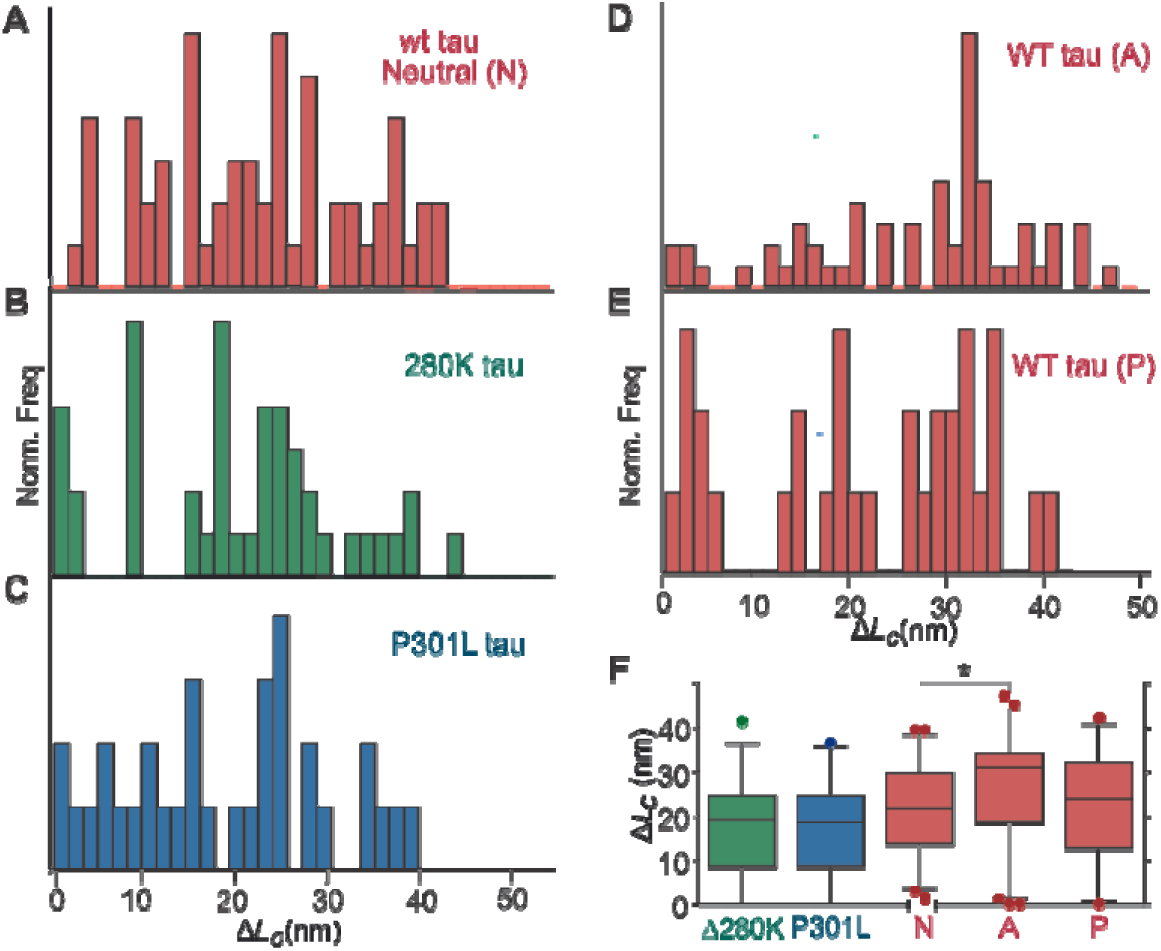
Mutations and buffer effects in the Δ*L*_*c*_ of the first mechanical barrier of tau. Δ*L*_*c*_ histograms of tau harboring pathological mutations **(A, B, C)** or in the three selected buffers **(A, D, E)** show a highly heterogeneous population in all the scenarios. **F)** Box plots with outliers for comparative analyses are shown. No statistical differences are observed among the structure sizes formed in conditions that promote the amyloidogenesis. A slight perturbation is quantified between the conformations acquired in neutral and antiamylodogenic conditions.

In a nutshell, buffer conditions and single-point mutations that exacerbate the amyloid aggregation of tau increase the frequency of NM conformers. In addition, buffer conditions, though no mutations, alter the types of conformations acquired by tau.

## Conclusion

The tau K18 fragment has been analyzed by AFM-SMFS using a mechanical protection strategy for unequivocal single-molecule identification. As it has been described for both pathological and functional aIDPs, monomeric tau shows a non-preferential rich conformational behavior, which includes hyper-mechanostable conformers (Hervas et al. 2012; 2016). Familial-disease mutations and a pro-amyloidogenic buffer increase the frequency of the unstructured NM population, which indicates that tau nucleation occurs in later steps **(Figure S5)**. A sub-population of the molecules remains heterogeneously structured, suggesting a source for polymorphic amyloid nuclei that can be altered by specific buffer conditions, although not by mutations. Our results open the door to a future nanomechanical screening of chemical libraries to select molecules able to stabilize the natural conformational polymorphism of tau, and thus, inhibit its amyloidogenesis.

## Materials and methods

### Molecular cloning

The coding region of the K18 fragment from tau was amplified by PCR using a plasmid clone containing the full length human tau as a template for PCR amplification, which was kindly provided by Prof. Jesús Ávila (CBMSO-UAM). The selected restriction sites used to insert the guest fragment into the carrier ubiquitin of the pFS-2 vector and the minimal fusion proteins were AgeI-MluI (Oroz, Hervas, and Carrion-Vazquez 2012; Fernandez-Ramirez et al. 2018). NheI-XhoI were selected for the insertion of K18 into pET28a (Novagen) expression vector. Mutants Δ280K and P301L were generated by site-directed mutagenesis using the *Phusion High Fidelity PCR kit*. The first cloning step was made in the pPCR2.1 (Invitrogen) vector and then the inset sequence was verified by sequencing both DNA strands. Primers were synthetized by Sigma-Aldrich **(Table 1)**.

**Table 1.**
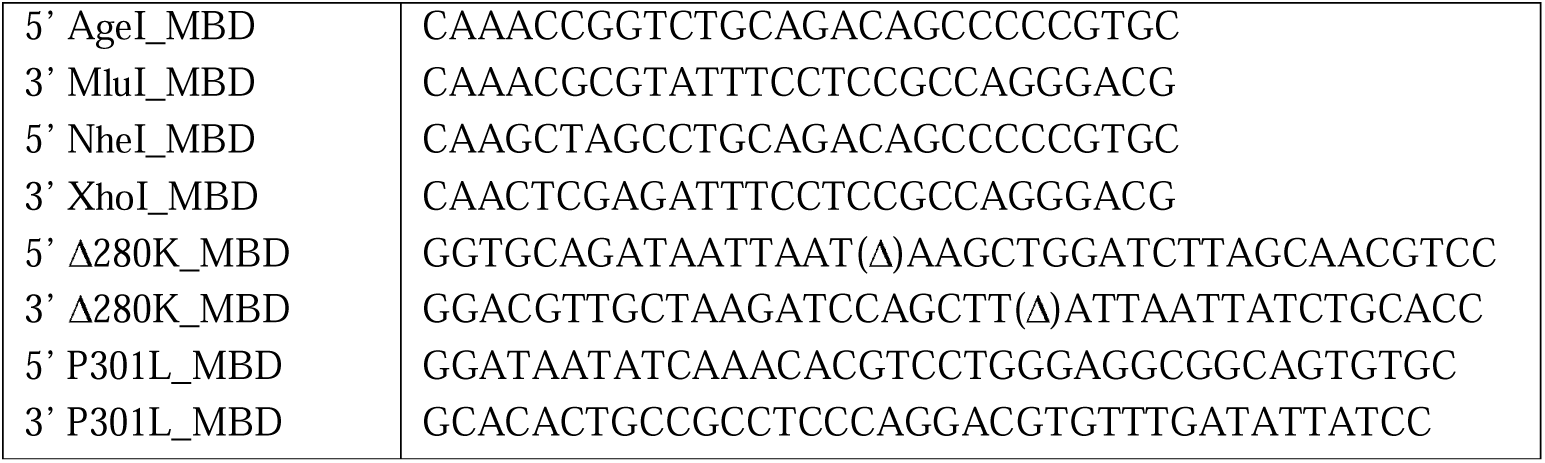
Primers used for the molecular cloning of tau inside the ubiquitin carrier, the expression vector and the insertion of single-point mutations.

### Protein expression and purification

Polyproteins, carrier-fused and isolated proteins were expressed in *E. coli* C41 (DE3) or BL21 (DE3) (Invitrogen) strains. The growth of bacterial cultures was done at 37°C and at 280-300 rpm stirring. When the OD_595_ reached 0.4-1, the protein expression was induced by 1mM IPTG for 4 h. The lysis of bacterial pellets was done by 1% Triton X-100, 0.5% Tween-20, sonication pulses and the addition of 1mg/ml lysozyme. The purification by Ni^2+^ affinity chromatography and size-exclusion was performed as previously described (Hervas et al. 2012; Fernandez-Ramirez et al. 2018), using an FPLC apparatus (ÄKTA Purifier, GE Healthcare). The concentration of proteins was determined by their absorbance at 280 nm using their molar extinction coefficients.

### DSC

Measurements were carried out at 50 *µ*M protein in 10 mM glycine/acetic buffer [pH 3.0] using a VP-DSC (MicroCal, LLC). Samples were scanned from 10 °C to 115 °C at a heating rate of 50 °C *per* hour, and the thermograms were analyzed using the DSC-Origin software after subtraction of the buffer *vs* buffer base line (Varea et al. 2000).

### Far-UV Circular dichroism (CD) spectroscopy

Measurements were carried out in a JASCO-J810 (JASCO Inc.) equipped with a Peltier temperature control unit. Samples were 5-20 *μ*M in 10 mM KH_2_PO_4_ [pH 7.0] and measured using quartz cuvettes of 1 mm pathlength. The corrected spectra were obtained by the subtraction of the buffer contribution. Then, they were converted into molar ellipticity using the average molecular mass per residue and the Spectra Manager software (Jasco Inc.)

### NMR

NMR experiments were performed on a Bruker Avance 800 MHz spectrometer fitted with Z-gradients and a triple resonance cryoprobe at 25.0 °C and pH 4.7. 1D ^1^H and 2D ^1^H NOESY spectra were registered on protein constructs consisting of tau K18 WT, P301L, and Δ280K alone or inserted into the ubiquitin carrier protein. 1D ^1^H spectra were recorded with 24-32 scans. The following acquisition parameters were used for 2D ^1^H NOESY spectra: Sweep width 12 x 12 ppm, number of scans *per* increment in the indirect dimension = 64-72, matrix size = 2k x 256 or 512. NOESY spectra were processed using the program TOPSPIN 2.1 (Bruker Biospin). Sodium 4,4-dimethyl-4-silapentane-1-sulfonate (DSS), at a concentration of 0.10 mM, was added as the chemical shift reference. Based on the published NMR assignments of human ubiquitin (Weber, Brown, and Mueller 1987), which were obtained at pH 4.7 and are deposited as BMRB entry 68, ^1^H signals characteristic of ubiquitin folded in its native conformation were identified and assigned in our spectra.

### TEM

20 *μ*M tau and Ubi-tau samples were incubated for 7 days at 37 °C and no stirring, in PBS [pH 7.5], 5 mM DTT and 0.02% NaN_3_, in presence or absence of 100 *μ*M QBP1. Carbon-coated 300-mesh copper grids were glow-discharged by an Emitech K100X apparatus (Quorum Technologies) and then, 10 *μ*L of each sample were adsorbed onto them. Samples were negatively stained with the addition of 1% uranyl acetate and, finally, cleaned by immersion in Milli-Q water.

### Dot Blot assay

After 7 days of incubation, 2 *μ*l of the tau and Ubi-tau samples incubated at 37 °C were spotted onto a nitrocellulose membrane. The membrane was blocked for 1 hour at room temperature with 10% non-fat milk in 0.01% Tween-20 TBS (TBS-T). Then, it was incubated with the fibril-specific, conformational monoclonal antibody OC (Millipore), prepared at a 1:1000 dilution in 3% BSA TBS-T. Then, the membrane was washed 3 times for 5 minutes using TBS-T before the next incubation for 1 hour with an anti-rabbit HRP conjugated secondary antibody [GE Healthcare] diluted 1:5000 in 3% BSA TBS-T. Finally, a new set of three washes with TBS-T was performed, and then the membrane was developed with ECL Plus chemiluminescence kit from Amersham-Pharmacia (GE Healthcare). Fibrillar species from A*β*_42_ were used as positive control.

### Single-cell protein microinjection

Microinjection was performed in COS-7 cell line cultured in 10% (v/v) fetal bovine serum (Life technologies) DMEM at 1·10^5^ cell/well density. Recombinant proteins at 2.5 *μ*M in PBS 7.4 were incubated for 7 days at 37 °C and then, each sample was double-blinded microinjected (n=3) with fluorescein-labeled dextran (Life Technologies) in 100-200 cells, using a micromanipulatior (Narishige). Ubiquitin, Ubi-MCS and VAMP2 were microinjected as negative controls. Cell viability was monitored by analyzing the fluorescein-positive cells and using a fluorescence microscope IX70 (Olympus), and the micrographs were acquired with a CCD camera (model C4742-95-12E3, Hamamatsu Photonics). 100% of cell viability was established 3 hours after the microinjection and the progression was monitored every 24 hours. The survival rate after 24 hours of microinjection was calculated by One-way ANOVA and *post-hoc* Tukey test. Viability data are presented as mean ± SEM.

### Single-molecule force spectroscopy

Ted Pella glass slides were functionalized by Nitrilotriacetic acid (NTA)-Ni^2+^ for the attachment of the fusion protein generated by the pFS-2 vector to the substrate by its His-tag. The cantilever (MLCT-AUNM, Veeco Metrology Group; and Biolever, Olympus) used was cleaned for ∼5 min with a UV lamp (UV/Ozone ProCleaner™ Plus, Bioforce Nanosciences Inc.) before each SMFS session, and the spring constant of each tip was calculated by applying the equipartition theorem (Florin et al. 1995) to the power spectrum.

Polyproteins were at 2-10 *μ*M in antiamyloidogenic, neutral or proamyloidogenic buffer and maintained at 4 °C between sessions. 3 molar excess of heparin and QBP1 20 *μ*M were added 15 minutes and the night before the SMFS experiments, respectively, and the protein was kept at 4 °C. Measurements were performed on a modified version of a home-made AFM instrument (Schlierf, Li, and Fernandez 2004; Valbuena et al. 2007) and in a commercial AFS instrument (Luigs & Neumann), at a constant pulling speed of 0.4 nm ms^-1^ in the length-clamp mode (Carrión-Vázquez et al. 2007). The stretched proteins were analyzed using Igor Pro 6 (Wavemetrics) and applying the WLC model of polymer elasticity (Bustamante et al. 2004):

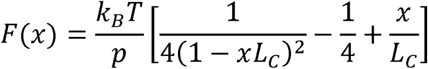

*L*_*C*_ (contour length of the stretched protein) and *p* (persistence length) are adjustable parameters, while the *F* refers to the stretching force and *x* refers to end-to-end distance.

#### Analysis of SMFS data

Very stringent criteria have been followed to identify single molecules in the SMFS recordings from the K18 (Hervas et al. 2012). First, the presence of a sawtooth pattern corresponding to the unfolding of ubiquitin molecular markers is required. We do not require a minimum number of unfolded markers since the carrier-guest strategy used in this study ensures the stretching of tau in an N-C direction, as is hosted in the ubiquitin carrier (Oroz, Hervas, and Carrion-Vazquez 2012). Second, the unfolding signal of the ubiquitin carrier must precede that from tau. However, it must be noted that the signal can appear either contiguously or interspersed (*i.e*., interrupted by *wt* ubiquitin markers). Third, considering the common presence of unspecific interactions in the proximal area of the AFM, unless the recording was completely noisy-free, the tau signal must appear after the first 50-70 nm. Fourth, after the unfolding of the ubiquitin carrier, distances not assigned to monomolecular markers must be of about ≈ 76.4 nm. This value represents the length of the carrier (≈26 nm) (Oroz, Hervas, and Carrion-Vazquez 2012) plus 50.4 nm from the 126-redidue K18 (Fernandez-Ramirez et al. 2018), considering 0.4 nm/residue (Ainavarapu et al. 2007).

#### Statistical analysis in SMFS experiments

χ-square test was used to analyze whether NM/M frequencies were significantly altered by the selected modulators (mutants, buffer conditions or QBP1 presence). The dependent variable “structure formation” can acquire NM or M values, and we analyzed its relationship with the independent variables “QBP1”, “buffer” and “mutations”. In testing the variable “QBP1”, we applied the Yates’ correction for continuity due to the small sample size and the one degree of freedom in the contingency table (Yates 1934).

## ACKOWLEDGEMENTS

This study was supported by the grants SAF2016-76678-C2-1-R (MC-V) and SAF2016-76678-C2-2-R (DVL) from the Spanish Ministry of Economy and Competitiveness as well as a grant of EU Joint Programme in Neurodegenerative Diseases (Spanish funding from the AC14/00037 grant, by the ISCIII) to MC-V. and DVL and grants of the Ministry of Economy and Competitiveness (BFU2015-70072-R) and the CIBER de Enfermedades Respiratorias (CIBERES; ISCIII) to MM. NMR experiments were performed in the “Manuel Rico” NMR laboratory (LMR) of the Spanish National Research Council (CSIC), a node of the Spanish Large-Scale National Facility (ICTS R-LRB).

## Notes

### Competing Interest Statement

The authors have declared no competing interest.

