## Supplemental Material for "Tau amyloidogenesis begins with a loss of its conformational polymorphism"

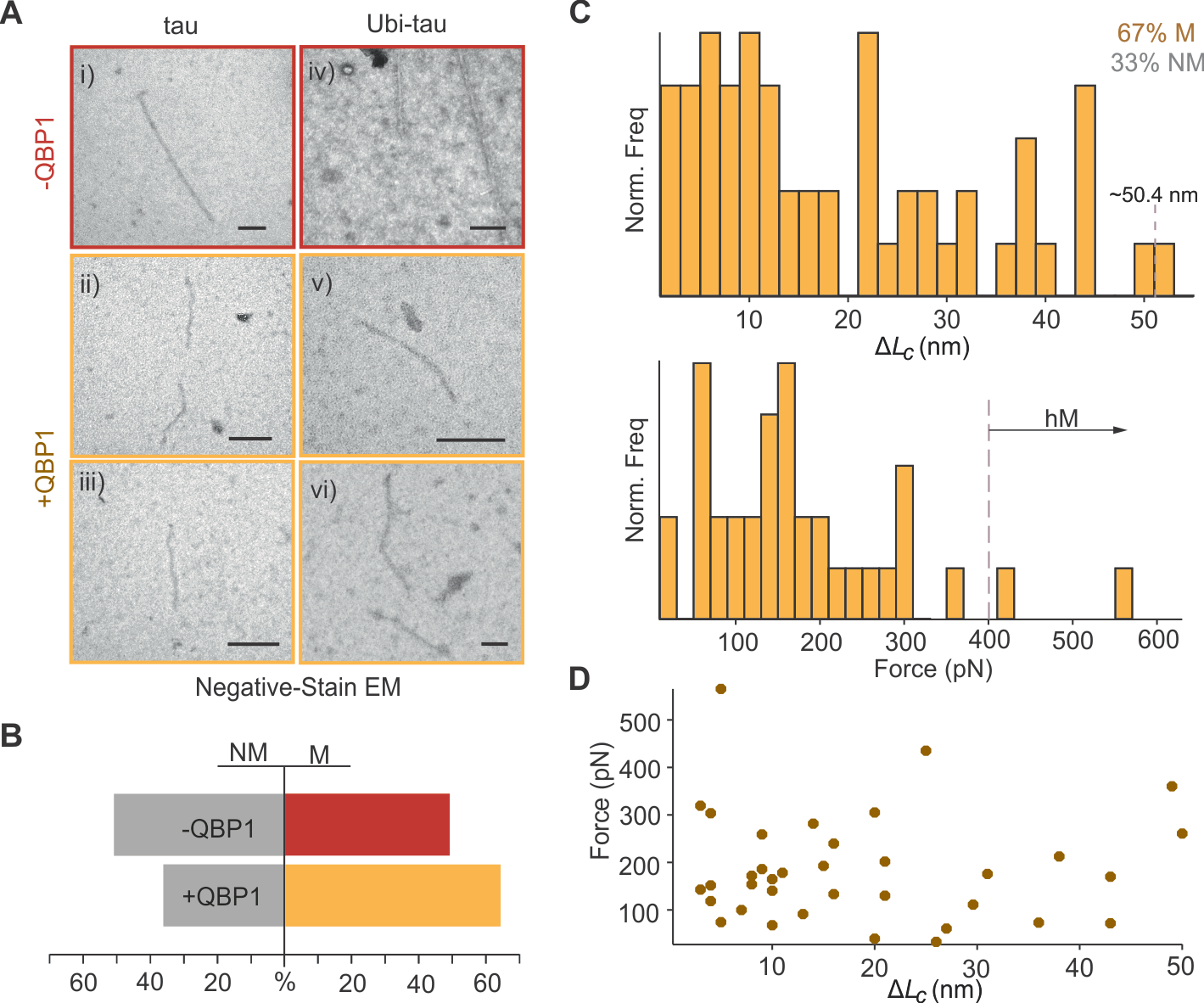

**Figure S1. Effect of QBP1 on tau. A)** Representative EM micrographs of tau fibers formed after 7 days of incubation. QBP1 does not prevent fiber formation of tau or Ubi-tau. Scale bar: 100 nm. **B)** Modulation tau nanomechanics by QBP1. In the presence of QBP1, 67% of the selected events were M (n=33). χ^2^ test of independence shows that QBP1 does not perturb the NM/M ratio (p>0.30) **C)** Histograms for Δ*L_c_* (n=57) and *F*_u_ (n=36) show a similar distribution as in the absence of QBP1 (see Figure 3). **D)** The scatter plot of Δ*L_c_* and *F*_u_ does not show the absence of specific classes of conformers.

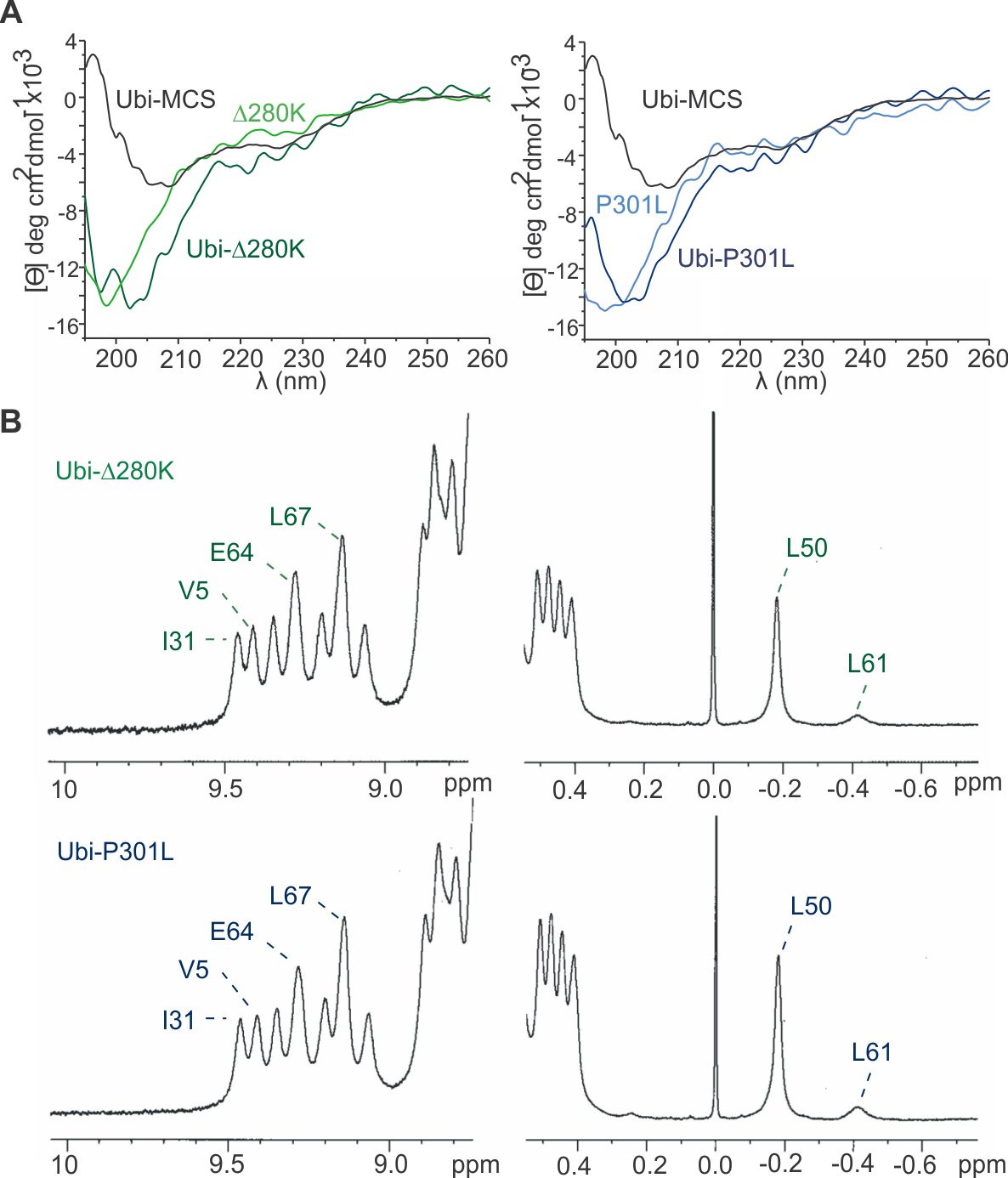

**Figure S2. Structural integrity of Ubi-*Δ*280K and Ubi-P301L A)** Far UV CD spectra of the fusion proteins show the contribution of Ubi-MCS and *Δ*280K or P301L signals, suggesting that both mutants maintain their features upon Ubi-MCS insertion. **B)** Magnified views of the ^1^HN (left) and upfield (right) regions of the ^1^H NMR spectra of Ubi-*Δ*280K and Ubi-P301L which show signals characteristic of folded ubiquitin.

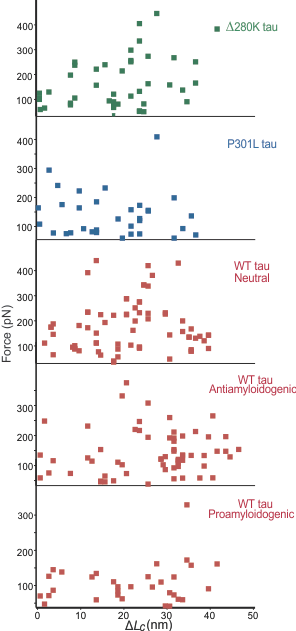

**Figure S3. Relation between Δ*L_c_* and *F*_u_ for the first mechanical barrier in different scenarios.** Scatter plots do not reveal any pattern, cluster or preferential conformation in any scenario studied.

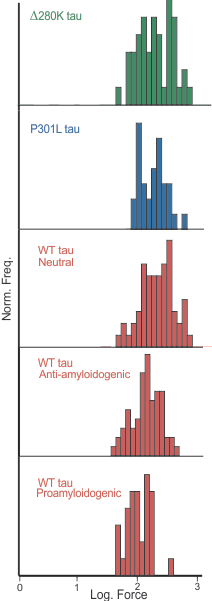

**Figure S4. *F*_u_ values transformed to a log-normal distribution.** Logarithm of *F*_u_ values obtained from the first mechanical barrier produce Gaussian distributions that pass the normality tests, and can be analyzed by parametric tests.

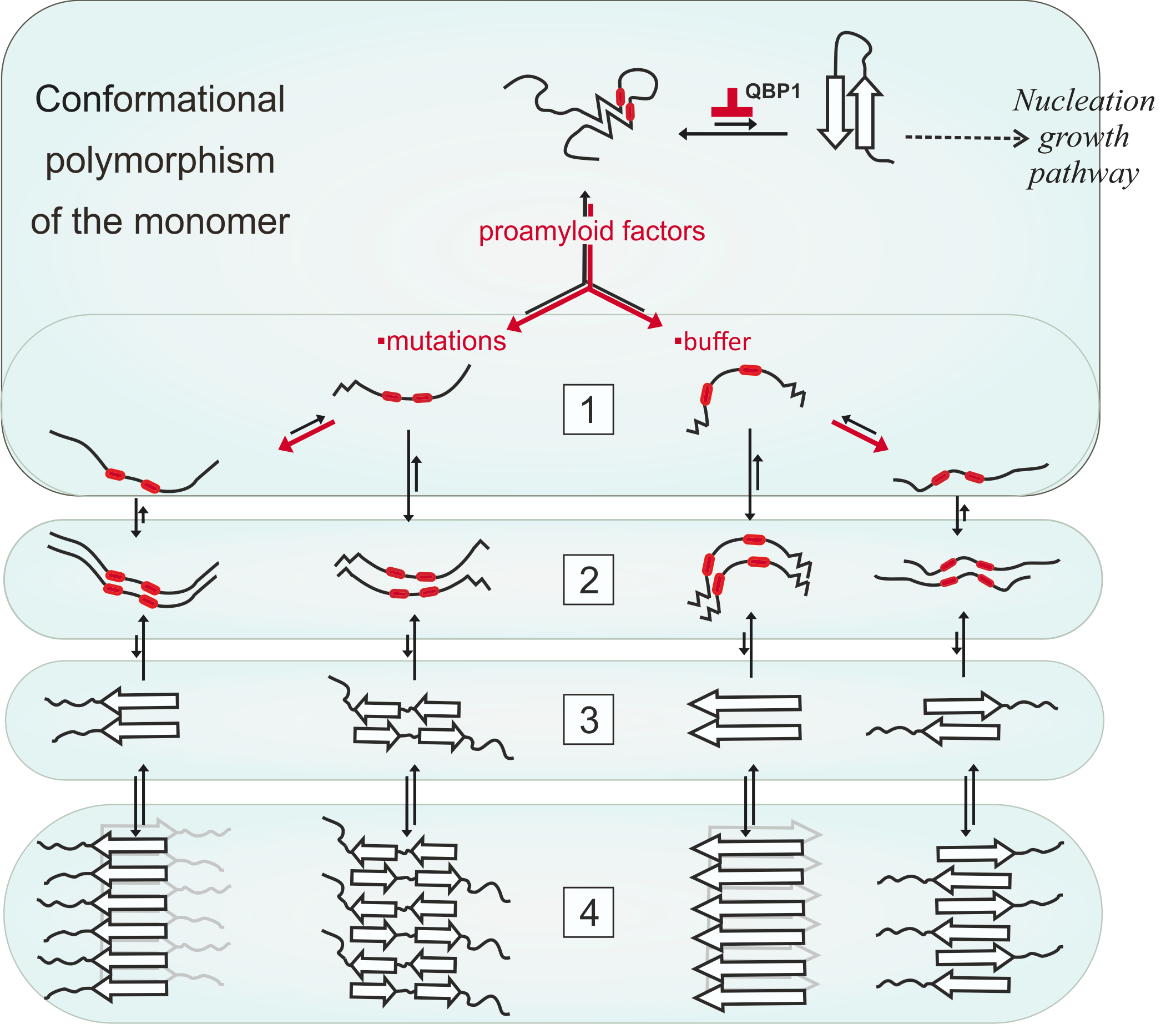

**Figure S5. Proposed mechanism for tau amyloid assembly.** Monomeric tau fluctuates among several conformations which are in a thermodynamic equilibrium, from NM random coils to highly M *β*-structured conformers. The conformational change that triggers the nucleation growth pathway is not favored. 1) Proamyloidogenic factors, like specific buffer conditions or pathological mutations favor an expanded tau conformation, in which the two pro-amyloid hexapeptides are exposed, corresponding with the amyloid-competent monomer. Intermolecular interactions driven by the aggregation-prone segments lead to the formation of early molten aggregates (2), which will later experience a structural rearrangement generating different aggregation nuclei (3), which evolve to form fibers with diverse morphologies (4).
